# Dynamics-driven allostery underlies pre-activation of the regulatory Ca^2+^-ATPase/phospholamban complex

**DOI:** 10.1101/2020.04.26.062299

**Authors:** Olga N. Raguimova, Rodrigo Aguayo-Ortiz, Seth L. Robia, L. Michel Espinoza-Fonseca

## Abstract

Sarcoplasmic reticulum (SR) Ca^2+^-ATPase (SERCA) and phospholamban (PLB) are essential for intracellular Ca^2+^ transport in myocytes. Ca^2+^-dependent activation of SERCA–PLB provides a rheostat function that regulates cytosolic and SR Ca^2+^ levels. While experimental and computational studies alone have led to a greater insight into the mechanisms for SERCA–PLB regulation, the structural changes induced by Ca^2+^ binding and how those are communicated to couple enzymatic activity with active transport remain poorly understood. Therefore, we have performed atomistic simulations totaling 32.7 μs and cell-based intramolecular fluorescence resonance energy transfer (FRET) experiments to determine structural changes of PLB-bound SERCA in response to Ca^2+^ binding. Complementary simulations and experiments showed structural disorder underlies PLB inhibition of SERCA, and Ca^2+^ binding is sufficient to shift the protein population toward a structurally ordered state of the complex. This structural transition results in a redistribution of structural states toward a partially closed conformation of SERCA’s cytosolic headpiece. Closure is accompanied by functional interactions between the N-domain β5-β6 loop and the A-domain. Regulation of these key structural elements indicate that Ca^2+^ is a critical mediator of allosteric signaling that dictates structural changes and motions that pre-activate SERCA–PLB. These findings provide direct support that dynamically driven protein allostery underlies PLB regulation of SERCA. These functional insights at unprecedented spatiotemporal resolution suggest a general modular architecture mechanism for dynamic regulation of the SERCA–PLB complex. Understanding these mechanisms is of paramount importance to guide therapeutic modulation of SERCA and other evolutionarily related ion-motive ATPases.

---

Sarcoplasmic Reticulum (SR) Ca^2+^-ATPase (SERCA) plays an essential role in normal cardiac function, clearing cytosolic Ca^2+^ during the diastolic phase of the cardiac cycle, relaxing muscle cells and allowing the ventricles to fill with blood [1]. SERCA pumps two Ca^2+^ ions into the SR lumen using energy derived from hydrolysis of ATP [2, 3] and is regulated by the 52-residue membrane protein phospholamban (PLB). PLB regulates SERCA by inhibiting its ability to transport Ca^2+^ [4, 5], and this inhibition is relieved by phosphorylation of PLB [6-10]. In the absence of PLB phosphorylation, binding of Ca^2+^ can still occur (though with lower apparent affinity), and this also leads to activation of the SERCA–PLB complex [11, 12]. Several regions and interactions have been shown to be essential for linking Ca^2+^ binding and SERCA–PLB reactivation, including (i) the N-terminal phosphorylation domain of PLB, which controls SERCA activity via order-to-disorder structural transitions [13-16], (ii) structural preorganization of the Ca^2+^ transport sites that facilitates SERCA activation [12, 17-19]; and (iii) the disruption of key inhibitory contacts involving the transmembrane domain of SERCA and the conserved PLB residue Asn34, a well-established regulatory element [20, 21].

A major challenge in molecular physiology and drug discovery is to understand the mechanisms of SERCA–PLB regulation in terms of the interactions and molecular motions of the underlying proteins. While the application of a variety of simulation and experimental approaches alone has led to a greater insight into the mechanisms for SERCA– PLB regulation, the structural changes induced by Ca^2+^ and how those are communicated to couple enzymatic activity with active transport remain poorly defined. More specifically, it remains unknown whether binding of a single Ca^2+^ ion initiates SERCA–PLB activation by stabilizing the transport sites [17, 22, 23] and producing a SERCA structure that is compatible with Ca^2+^ binding [24], or by inducing large-scale domain arrangements that produce catalytically competent conformations of SERCA [25, 26]. Answering these questions requires knowledge of the structural changes and motions at high spatiotemporal resolution. Therefore, we have used long, independent molecular dynamics (MD) simulations complemented with cell-based intramolecular fluorescence resonance energy transfer (FRET) experiments to identify the fundamental mechanisms for Ca^2+^-mediated activation of the SERCA–PLB regulatory complex. The result is a vivid atomic-resolution visualization of the dynamic regulation of the Ca^2+^ transport machinery in the heart.

We performed atomistic MD simulations totaling 32.7 μs (see Methods, *Supporting Information*) to identify the conformational changes induced by Ca^2+^ in SERCA bound to unphosphorylated PLB. Root-mean-square deviation analysis showed that the complex is structurally in the presence and absence of bound Ca^2+^ (**Fig. S1–S9**, *Supporting information*). Ca^2+^ binding to the complex does not induce changes in the secondary structure of PLB or the occupancy fraction of PLB in the canonical binding groove. However, binding of Ca^2+^ disrupts key inhibitory contacts between residues Asn34 of PLB and Gly801 of SERCA (**Fig. S1–S9**, *Supporting information*). These findings agree with previous studies showing that binding of a single Ca^2+^ ion relieves inhibitory SERCA–PLB contacts without inducing changes in the overall architecture of the SERCA–PLB interface[17] or secondary structure of PLB, such as those induced by PLB phosphorylation [13-16, 27].

We analyzed the trajectories using Cartesian principal component analysis (cPCA) to systematically reduce the number of dimensions needed to describe protein dynamics while retaining the essential space [28]. Atomic fluctuation of SERCA in the essential space defined by the first 5 eigenvectors (**Fig. S10**, *Supporting information*), showed that binding of Ca^2+^ to the transmembrane transport sites (**Fig. 1**) decreases the amplitude of motion in the ion transport domain [29, 30], namely in transmembrane helices M1, M3 and M4, as evidenced by a decrease in RMSF for this region (**Fig. 1**). These results correlate well with reports showing that helices M1, M3 and M4 must move to cluster together and form the transport sites [31], with Ca^2+^ binding facilitating this process by reducing the mobility of these helices. Substantial structural changes are also observed in the luminal side of SERCA, mainly in the large loop that connects transmembrane helices M7 and M8 (loop M7-M8, principal components 1-2 and 4-5). This finding is significant because loop M7-M8 has been proposed to stabilize the luminal pathway required for Ca^2+^ translocation[32] as well as for ion conduction in other P-type pumps (e.g., Na^+^-K^+^-ATPase) [33].

**Fig. 1.**
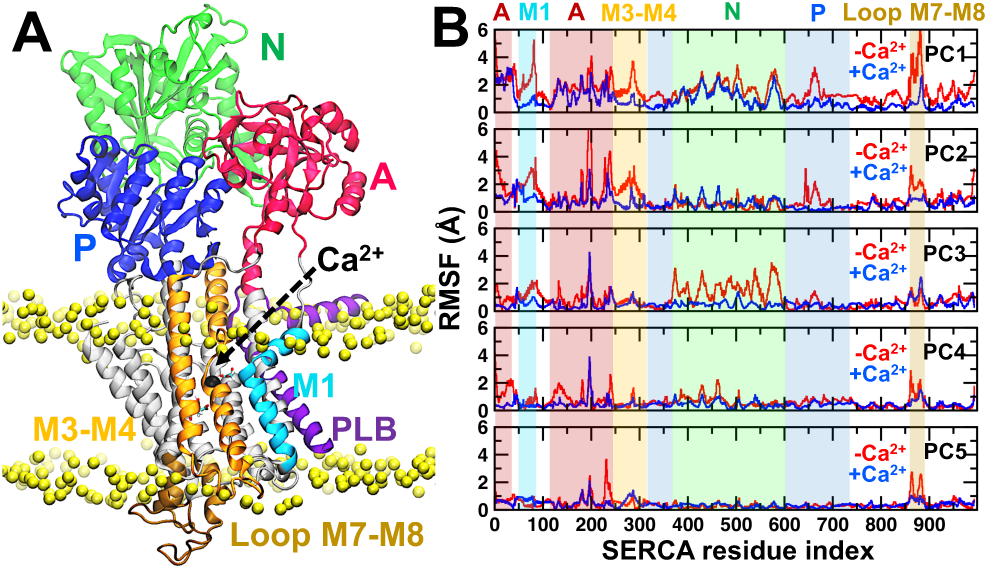
Comparison of domain flexibility between Ca^2+^-free and Ca^2+^-bound SERCA–PLB. (A) Structure of the SERCA–PLB (ribbons) complex embedded in a POPC lipid bilayer (yellow spheres). PLB is shown in purple, and SERCA is colored according to its four functional domains: N (green), P (blue), A (red), and TM (gray); additionally, helices M1, M3-M4 and the luminal loop M7-M8 are shown in cyan, orange and brown, respectively. Ca^2+^ bound in the transport site I is shown as a black sphere. (B) Cα root-mean square fluctuation (RMSF) of SERCA in the essential space defined by the first 5 eigenvectors calculated using the combined MD trajectories of Ca^2+^-free (red) and Ca^2+^-bound SERCA–PLB. The shaded areas indicate the location of functionally important domains in the sequence of the protein shown in panel A.

The effects of Ca^2+^ binding also propagate to the cytosolic headpiece of the protein (**Fig. 1**), primarily affecting the dynamics of the actuator (A) domain (PC1 through PC5), residues 640-670 of the phosphorylation (P) domain (PC1 and PC2), and the nucleotide-binding (N) domain (PC3). To understand these Ca^2+^-induced structural shifts in terms of domain motions, we obtained cPCA-based motion representation of SERCA obtained using the Ca^2+^-bound and Ca^2+^-free complexes (**Fig. 2**). In the absence of Ca^2+^, the relevant motions of the cytosolic headpiece of SERCA are a scissor-like motion of the N and A/P domains, with a movement similar to that of scissor blades opening (PC1, **Fig. 2A**), and a counterclockwise rotation of the N and A domains, with a movement similar to that of a rotor moving about the membrane normal axis (PC3, **Fig. 2A**). Conversely, Ca^2+^ binding to the complex induces counterrotation of the N and A domains about the membrane normal axis (PC1, **Fig. 2B**), a counterclockwise rotation of the N domain (PC2, **Fig. 2B**) and the displacement of the N and A domains in opposite directions, similar to that of the ‘opening’ of the headpiece (PC3, **Fig. 2B**). The headpiece motions captured by the MD simulations are consistent with the structural transitions of SERCA in the absence of PLB, where apo SERCA populates a partially open cytosolic headpiece prior to Ca^2+^-induced activation [25, 34], with SERCA undergoing a structural shift toward a closed conformation during Ca^2+^ signaling and activation [25, 35].

**Fig. 2.**
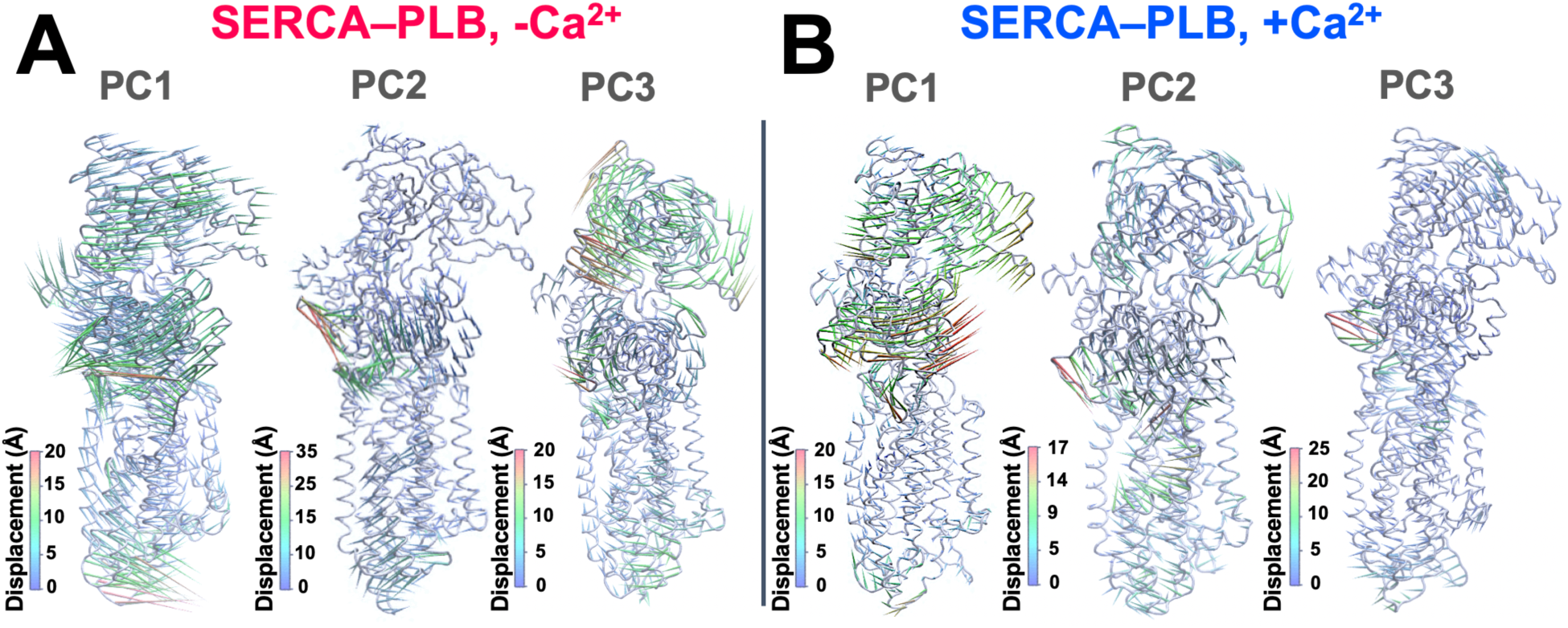
Porcupine representation of the domain motions described by the first three principal components. (A) Porcupine plot motions of SERCA detected from MD ensembles of SERCA–PLB in the absence of bound Ca^2+^. (B) Changes in domain motions of SERCA induced by binding of a single Ca^2+^ to the SERCA–PLB complex. In all cases, the SERCA shown as a cartoon, and the arrows indicate the direction of rigid-body motion and colored based on the atomic displacement of the Cα atoms.

cPCA-based structural landscape showed that while Ca^2+^-free SERCA–PLB samples structures that are similar to that of crystal structure (PDB: 4ky t[36], **Fig. 3A**), the structural population of the complex is heterogenous in solution, thus suggesting that the crystal structure represents only a only a small portion of the overall complex conformation. Ca^2+^ binding to the transport sites produces a structurally more homogeneous ensemble that is more consistent with the crystal structure of the complex (PDB: 4kyt [36]), but also populates a conformation of the complex that resemble that of a compact and structurally ordered state of SERCA (PDB:1vfp [37], **Fig. 3B**). The differences in structural landscape and domain motions captured by our µs-long simulations suggest that Ca^2+^ binding to SERCA–PLB induces a disorder-to-order transition in the headpiece of SERCA. We tested this hypothesis by quantifying intramolecular FRET between fluorescent proteins fused to the A- and N-domains of SERCA (2-color SERCA) as an index of the structural changes of the SERCA cytoplasmic headpiece [25, 35, 38, 39]. We explored the Ca^2+^-dependent change in FRET of microsomal preparations of 2-color SERCA co-expressed with the non-phosphorylatable PLBS16A mutant to mimic the MD conditions, and 2-color SERCA alone as a control (see Methods, *Supporting Information*). We observed increased intramolecular FRET for 2-color SERCA co-expressed with PLBS16A (**Fig 3C**, blue trace) and FRET increased with increasing free Ca^2+^ (EC50 = 850 nM). This change is consistent with a direct effect on the overall shape of the SERCA cytoplasmic headpiece, as revealed by Ca^2+^-dependent intramolecular FRET of the 2-color SERCA alone (**Fig. 3C**, black trace). The FRET data are consistent with our MD-based model that Ca^2+^-binding stabilizes a SERCA–PLB regulatory complex characterized by a compact, ordered headpiece conformation. The agreement between complementary MD simulations and FRET experiments support the notion that structural disorder underlies PLB inhibition of SERCA, and Ca^2+^ binding is sufficient to shift the complex toward a compact structure of the headpiece.

**Fig. 3.**
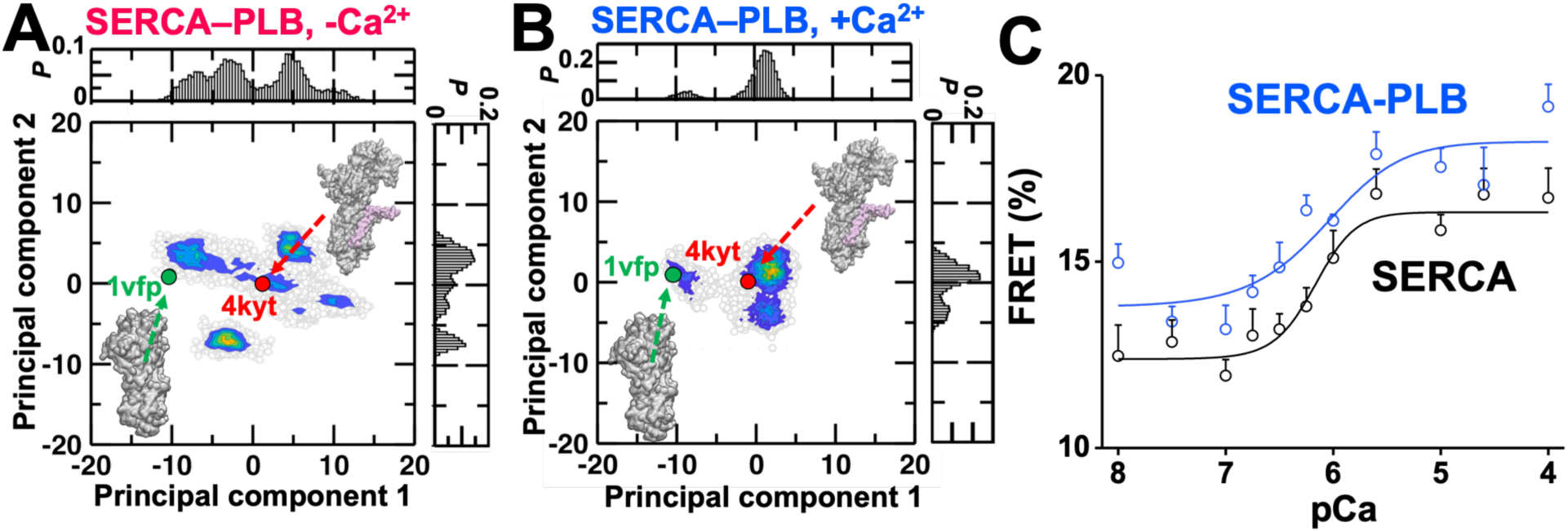
Differential structural dynamics of SERCA in response to Ca^2+^ binding determined by complementary MD simulations and FRET experiments. Structural landscape of SERCA–PLB in (A) the absence of Ca^2+^ and (B) the presence of a single Ca^2+^ ion bound in the transport sites of SERCA. For references, we show the location of the crystal structures of SERCA–PLB (4kyt, in red) and SERCA bound to two Ca^2+^ ions and AMP-PCP (1vfp, in green). (C) Change in FRET of 2-color SERCA in response to increase in Ca^2+^ concentrations as detected by confocal fluorescence microscopy. Ca^2+^ dependent FRET was measured for SERCA–PLB (blue trace) and SERCA alone as a control (black trace). Data are shown as mean ± SD (*n*=3).

Previous studies have shown headpiece misalignment can occur despite SERCA populating compact headpiece structures [26], so we asked whether Ca^2+^-induced changes observed in our MD simulations brings together key structural element required for SERCA activation. We focus on Ca-Ca distances between two sets of residues shown in **Fig. 4A**: Arg139 (A domain) and Asp426 (loop Nβ5-β6), which play a key role in initiating the open-to-closed conformational change of SERCA [25, 40], and Thr171 (A domain) and Glu486 (N domain), which come together to stabilize a compact headpiece conformation that precedes ATP hydrolysis and Ca^2+^ transport [37]. We tested a one-through four-Gaussian distributions for each set of Ca-Ca distances, and the best fit was selected based on *χ*^2^ values (**Tables S1– S2**, *Supporting Information*). In the absence of Ca^2+^, the distances Thr171–Glu486 and Arg139–Asp426 fit to four and two Gaussian subpopulations, respectively (**Fig. 4B**). These populations are consistent with an open and dynamically disordered SERCA headpiece because they cover a wide range of distances (i.e., distances between 10–45 Å) and deviate from the crystal structures of SERCA–PLB (PDB: 4kyt [36]) and the structurally ordered state of the pump (PDB: 1vfp [37], **Fig. 4B**). Conversely, Ca^2+^ binding induces a decrease in the N-A interdomain distances (**Fig. 4C**), thus facilitating the formation of a more dynamically ordered, compact structure of the cytosolic headpiece. Interestingly, the Ca^2+^-bound complex populates a relatively small conformation where Arg139 and Asp426 come close together similar to that found in the crystal structure of SERCA that precedes full activation (**Fig 4C**, right panel, green trace) [37]. This observation is significant because the interaction between the loop Nβ5-β6 and residues 133-139 of the A-domain has been proposed to be an important step that initiates SERCA activation [25, 40]. These specific contacts and concerted motions are suggestive of activation-precursor events, and also suggest that activation of the complex occurs as a series of sequential structural steps controlled primarily by Ca^2+^ binding [22, 41].

**Fig. 4.**
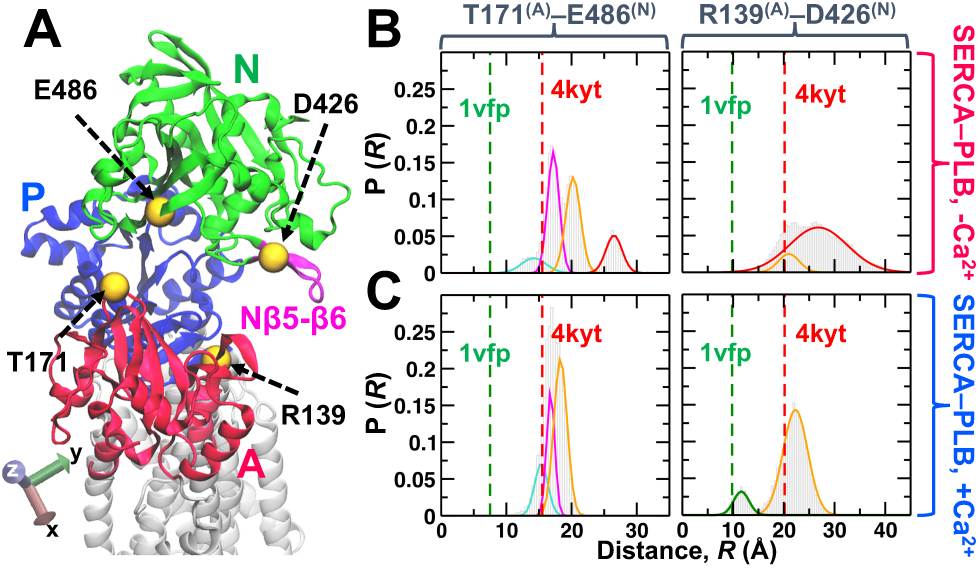
Structural arrangement of the A and N domains of SERCA. (A) Ribbon representation of SERCA showing the cytosolic N, A and P domains in green, red and blue, respectively. Functional loop Nβ5-β6 located in the N domain is shown in pink. Yellow spheres indicate the position of residues used to calculate interdomain distance distributions. Cα-Cα distance distributions between Thr171– Glu486 and Arg139–Asp426 obtained for the (B) Ca^2+^-free and the (C) Ca^2+^-bound SERCA–PLB complex. For comparison, discrete distances for the same pairs of residues were calculated from crystal structures of the apo SERCA–PLB complex (4kyt, in red) and SERCA bound to two Ca^2+^ ions and AMP-PCP (1vfp, in green).

We propose a mechanistic model that explains how the Ca^2+^-binding signal transmitted to the cytosolic region of SERCA. First, we note in the absence of Ca^2+^, PLB favors a disordered ‘off’ state of SERCA, where the structural arrangement required for Ca^2+^ occlusion and alignment of the catalytic elements is an infrequent event, thus rendering SERCA catalytically inactive. Ca^2+^ binding decreases the mobility of the transmembrane domain and allosterically induces concerted rigid-body motions of the cytosolic domains that bring loop Nβ5-β6 and residues Arg139 at the N-A domain interface (**Fig. 4A**). We propose that this step is a critical kinetic factor that precedes activation by (i) dictating the structural changes and motions that facilitate cooperative activation of SERCA following binding of the second Ca^2+^ ion [42-44], and (ii) favorably orienting the catalytic elements required for coupling of ATP hydrolysis subsequent Ca^2+^ transport [25, 26, 37, 40]. This mechanistic model suggests that binding of a single Ca^2+^ is a critical mediator of allosteric signaling because it controls key structural elements that are necessary to populate a pre-activated state of the SERCA– PLB complex.

In summary, we provide direct support that dynamically driven protein allostery [45] underlies Ca^2+^-mediated pre-activation of SERCA–PLB. Our findings suggest a general modular architecture mechanism for dynamic regulation of SERCA and other ion-motive ATPases, where modular domains within a single protein allows complex regulation while conserving the structure of the individual domains [46]. Here, regulation of SERCA is achieved by stringing together its catalytic elements with regulatory modules (e.g., loop Nβ5-β6) that serve as checkpoints during intracellular Ca^2+^ signaling [25, 40]. Ultimately, each module provides a combinatorial enhancement to its regulation and function in the cell [47]. Unraveling these modular architecture mechanisms at molecular level are of paramount importance to initiate mechanism-based design of therapies targeting SERCA and other evolutionarily related ion-motive ATPases [48, 49].

## SUPPORTING INFORMATION

Methods, Figures S1–S10, Tables S1-S2, and Supporting Information References.

## ACKNOWLEDGMENT

This work was supported by the National Institutes of Health grants R01GM120142 and R01HL148068 (to L.M.E.-F.), and R01HL092321 and R01HL143816 (to S.L.R.), and from the Loyola University Chicago Cardiovascular Research Institute (to S.L.R.). This research was supported in part through computational resources and services provided by Advanced Research Computing at the University of Michigan, Ann Arbor.\

## Notes

### Competing Interest Statement

The authors have declared no competing interest.

