## Supplemental Data 1 for "Dynamics-driven allostery underlies pre-activation of the regulatory Ca^2+^-ATPase/phospholamban complex"

### TABLE OF CONTENTS


| Contents | | Page |
|---|---|---|
| **Materials and Methods** | | S2 |
| **Fig. S1** | Structural analysis of the $Ca^{2+}$-free SERCA–PLB from the MD replicate #1. | S4 |
| **Fig. S2** | Structural analysis of the $Ca^{2+}$-free SERCA–PLB from the MD replicate #2. | S5 |
| **Fig. S3** | Structural analysis of the $Ca^{2+}$-free SERCA–PLB from the MD replicate #3. | S6 |
| **Fig. S4** | Structural analysis of the $Ca^{2+}$-free SERCA–PLB from the MD replicate #4. | S7 |
| **Fig. S5** | Structural analysis of the $Ca^{2+}$-free SERCA–PLB from the MD replicate #5. | S8 |
| **Fig. S6** | Structural analysis of the $Ca^{2+}$-bound SERCA–PLB from the MD replicate #1. | S9 |
| **Fig. S7** | Structural analysis of the $Ca^{2+}$-bound SERCA–PLB from the MD replicate #2. | S10 |
| **Fig. S8** | Structural analysis of the $Ca^{2+}$-bound SERCA–PLB from the MD replicate #3. | S11 |
| **Fig. S9** | Structural analysis of the $Ca^{2+}$-bound SERCA–PLB from the MD replicate #4. | S12 |
| **Fig. S10** | Probability distributions for the first 15 principal components extracted in the trajectories of $Ca^{2+}$-free and $Ca^{2+}$-bound SERCA–PLB complex. | S13 |
| **Table S1** | Correlation coefficient ($r$) and Pearson's test ($\chi^2$) values computed for the Gaussian fitting models applied to the distance distribution between residues Thr171 and Glu486 obtained from the trajectories of $Ca^{2+}$-free and $Ca^{2+}$-bound SERCA–PLB. | S14 |
| **Table S2** | Correlation coefficient ($r$) and Pearson's test ($\chi^2$) values computed for the Gaussian fitting models applied to the distance distribution between residues Arg139 and Asp426 obtained from the trajectories of $Ca^{2+}$-free and $Ca^{2+}$-bound SERCA–PLB. | S14 |
| **Supplementary References** | | S15 |

### Supporting Information

#### Materials and Methods

*Setting up the SERCA–PLB complex at free $Ca^{2+}$ conditions*

We used an atomic model of the full-length SERCA–PLB structure[1] to simulate the inhibited complex at free $Ca^{2+}$ conditions. On the basis of our previous studies [1], we modeled transport site residues Glu309 and Asp800 as unprotonated and residues Glu771 and Glu908 as protonated. In addition, we adjusted the $pK_a$ of other ionizable residues to a pH value of ~7.2 using PROPKA [2, 3]. The complex was inserted in a pre-equilibrated 120×120 Å bilayer of POPC lipids. We used the first layer phospholipids that surround SERCA in the E1 state [4] as a reference to insert the complex in the lipid bilayer. This initial system was solvated using TIP3P water molecules with a minimum margin of 15 Å between the protein and the edges of the periodic box in the z-axis. $K^+$, and $Cl^-$ ions were added to neutralize the system and to produce a KCl concentration of ~100 mM.

*Setting up the $Ca^{2+}$-bound SERCA–PLB complex*

We used the structure of the complex bound to a single $Ca^{2+}$ ion obtained previously in our group [5] as a starting structure to simulate the SERCA–PLB complex at saturating $Ca^{2+}$ conditions. For these simulations, we modeled transport site residues Glu309, Glu771 and Asp800 as unprotonated and residue Glu908 as protonated. The complex was inserted in a pre-equilibrated 120×120 Å bilayer of POPC lipids. The complex was solvated using TIP3P water molecules; the final lipid-water-protein complex was prepared using the same protocol and KCl concentrations used for the complex. Given the size of the system, we found that a single $Ca^{2+}$ ion bound to the transport sites of SERCA corresponds to a total $Ca^{2+}$ concentration of ~400 µM. This total $Ca^{2+}$ concentration is much higher than that estimated at rest [6], and is also in good agreement with previous estimates at elevated cytosolic $Ca^{2+}$ [7-9].

*Molecular dynamics simulations*

MD simulations of all systems were performed by using the program NAMD,[10] with periodic boundary conditions [11], particle mesh Ewald [12, 13], a non-bonded cutoff of 12 Å, and a 2 fs time step. CHARMM36 force field topologies and parameters were used for the proteins [14], lipid [15], water, $Ca^{2+}$, $K^+$ and $Cl^-$. The NPT ensemble was maintained with a Langevin thermostat (310K) and an anisotropic Langevin piston barostat (1 atm). Fully solvated systems were first subjected to energy minimization, followed by gradually warming up of the systems for 200 ps. This procedure was followed by 10 ns of equilibration with backbone atoms harmonically restrained using a force constant of 10 kcal $mol^{-1}$ $Å^{-2}$. We performed five independent MD simulations of the $Ca^{2+}$-free complex and four of the $Ca^{2+}$-bound SERCA–PLB system. Details of the simulation times are shown below:

| **$Ca^{2+}$-free SERCA–PLB** | | **$Ca^{2+}$-bound SERCA–PLB** | |
|---|---|---|---|
| MD replicate number | Simulation time (µs) | MD replicate number | Simulation time (µs) |
| Replicate #1 | 3.73 | Replicate #1 | 3.95 |
| Replicate #2 | 3.77 | Replicate #2 | 3.68 |
| Replicate #3 | 3.36 | Replicate #3 | 3.83 |
| Replicate #4 | 3.40 | Replicate #4 | 3.46 |
| Replicate #5 | 3.53 | | |

*Cartesian Principal component analysis*

We used Cartesian Principal Component Analysis (cPCA) to characterize the essential space of SERCA, and to determine whether the structural changes observed in our MD simulations are associated with stabilization or destabilization of the E2 state. cPCA uses the actual dynamics of the protein to generate the appropriate collective coordinates that capture the important structural and dynamic features of the native states of the protein [16]. For cPCA, we aligned structures using the 10-helix transmembrane domain of SERCA as a reference. We then projected the trajectories of the SERCA into the phase space; the projection of a trajectory on the eigenvectors of its covariance matrix is called principal component. We then constructed probability histograms of each principal component; each histogram is then fitted to a one-Gaussian distribution to determine whether a principal component belongs to essential phase space [16]. All cPCA calculations and analyses were performed using the GROMACS package [17].

*Analysis and visualization*

We calculated the backbone root mean square deviation (RMSD) for SERCA, cytosolic domains (CD) and transmembrane region (TM), and PLB, N-terminal domain (NT) and TM. PLB secondary structure (helicity) through the simulation was computed employing *dssp* program [18] integrated in the built-in tool *do_dssp*. We calculate the occupancy fraction (OF) of SERCA residues (center of mass) located within 3.0 Å of PLB and PLB residues (center of mass) located within 3.0 Å of SERCA using the *select* built-in tool implemented in GROMACS.[17] Hydrogen bonds were computed using *HydrogenBondAnalysis* library of MDAnalysis. Rendering of structures was performed with VMD [19].

### *Fluorescence resonance energy transfer (FRET) experments*

Endoplasmic reticulum (ER) microsomal membranes of 2-color SERCA in the presence or absence of phosphomimetic PLB were prepared as described previously [20]. Briefly, HEK-293 cells were infected with adenoviruses encoding non-fluorescent PLB and/or 2-color SERCA with cerulean fused to the N-terminus and yellow fluorescent protein (YFP) inserted before residue 509 [21]. Cells were homogenized in a solution of 0.5 mM $MgCl_2$, 10 mM Tris-HCl, pH 7.5 containing protease inhibitor cocktail using a Potter-Elvehjem homogenizer followed by repeated passage through a 27-gauge needle. ER microsomal membranes were obtained from cell homogenates by successive centrifugation steps at 1,000 x g and 126,000 x g. Microsomes were diluted in a Ca-free base solution of 100 mM KCl, 5 mM $MgCl_2$, 2 mM EGTA, 10 mM imidazole, pH 7.0 and $CaCl_2$ was added to obtain a range of free $Ca^{2+}$ concentrations to span pCa 8 to pCa 4 [22]. Microsomes were pipetted onto glass coverslips and FRET was quantified by acceptor-sensitization using widefield fluorescence microscopy as described previously [20].

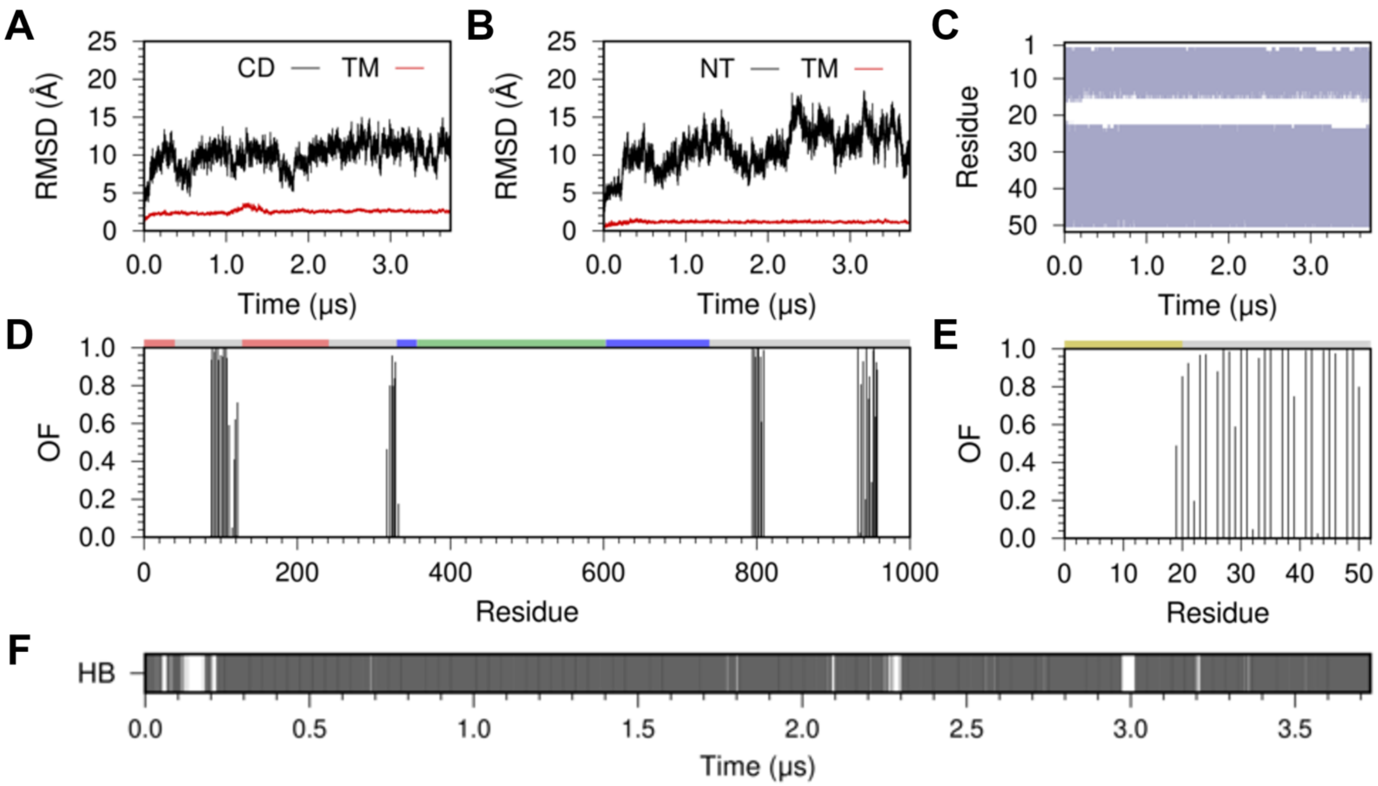


**Fig. S1. Structural analysis of the $Ca^{2+}$-free SERCA–PLB from the MD replicate #1**. (A) The RMSD of the cytosolic (CD, black) and transmembrane (TM, red) domains of SERCA. (B) The RMSD of the cytosolic N-terminal (NT, black) and transmembrane (TM, red) regions of PLB in the complex. In all cases, the RMSD was calculated by aligning the backbone of the TM domain of SERCA with the structure at the beginning of the MD replicate. (C) Secondary structure evolution of PLB in the complex; here, α-helix and random coil are colored as violet and white, respectively. This plot shows that the structure of PLB remains intact in the μs timescale. (D) Occupancy fraction (OF) of PLB in the canonical binding site of SERCA calculated from the MD replicate. This plot indicates that PLB interacts primarily with the TM domain of SERCA (gray) and does not form favorable interactions with the A (red), P (blue) or N (green) domains of SERCA. (E) Occupancy fraction (OF) of each individual residue of PLB in the canonical site. This plot demonstrates that the TM (gray), but not the cytosolic (yellow) domain of PLB interacts with SERCA in the MD trajectory. (F) Time-dependent evolution of the hydrogen bond between PLB residue Asn34 and SERCA residue Gly801. High stability of this interaction is key for SERCA inhibition by PLB [1, 5, 23], thus demonstrating that in the absence of $Ca^{2+}$ SERCA is primarily in an inhibited state.

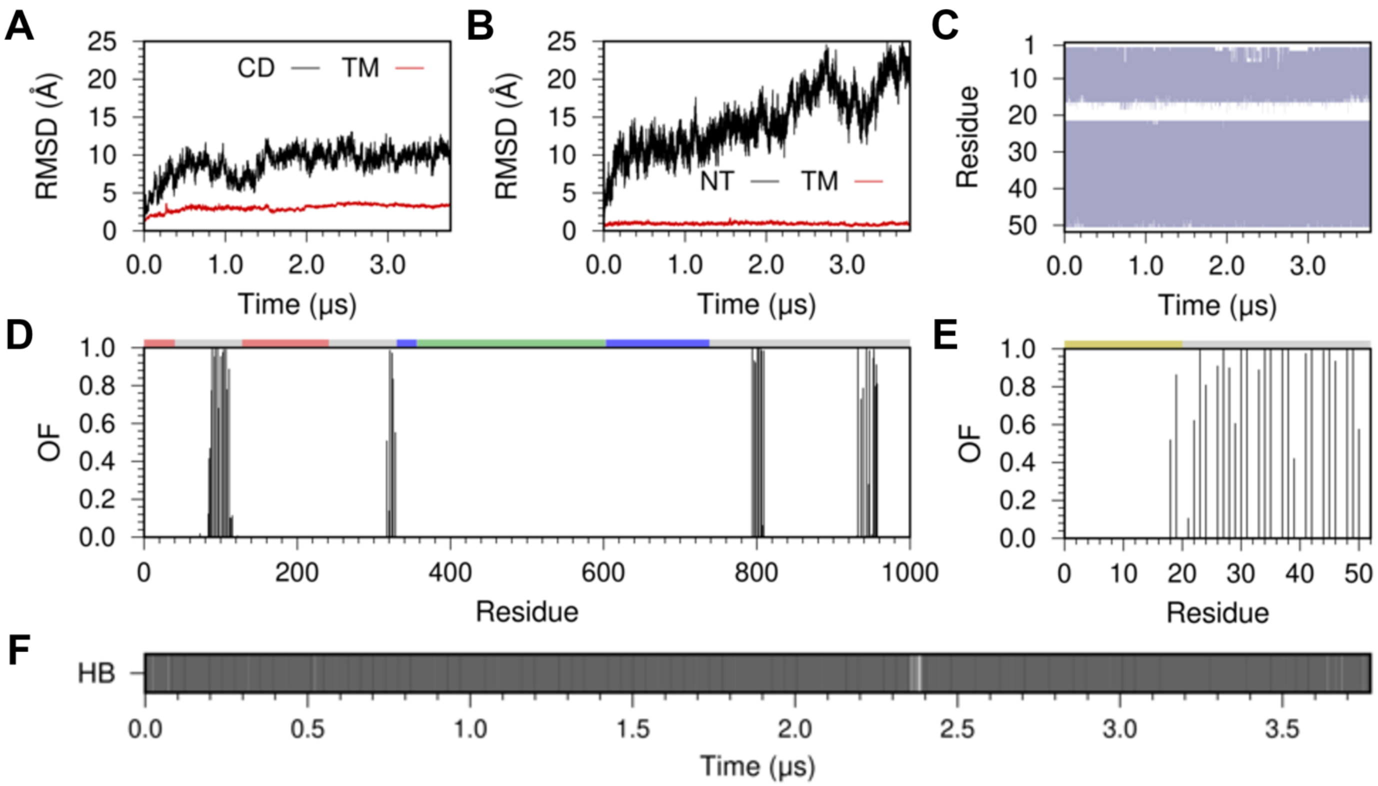


**Fig. S2. Structural analysis of the $Ca^{2+}$-free SERCA–PLB from the MD replicate #2**. (A) The RMSD of the cytosolic (CD, black) and transmembrane (TM, red) domains of SERCA. (B) The RMSD of the cytosolic N-terminal (NT, black) and transmembrane (TM, red) regions of PLB in the complex. In all cases, the RMSD was calculated by aligning the backbone of the TM domain of SERCA with the structure at the beginning of the MD replicate. (C) Secondary structure evolution of PLB in the complex; here, α-helix and random coil are colored as violet and white, respectively. This plot shows that the structure of PLB remains intact in the μs timescale. (D) Occupancy fraction (OF) of PLB in the canonical binding site of SERCA calculated from the MD replicate. This plot indicates that PLB interacts only with the TM domain of SERCA (gray) and does not form interactions with the A (red), P (blue) or N (green) domains of SERCA. (E) Occupancy fraction (OF) of each individual residue of PLB in the canonical site. This plot demonstrates that the TM (gray), and some residues in the cytosolic extension (residues Ile18 and Glu19) domain of PLB interacts with SERCA in the MD trajectory. (F) Time-dependent evolution of the hydrogen bond between PLB residue Asn34 and SERCA residue Gly801. High stability of this interaction is key for SERCA inhibition by PLB [1, 5, 23], thus demonstrating that in the absence of $Ca^{2+}$ SERCA is primarily in an inhibited state.

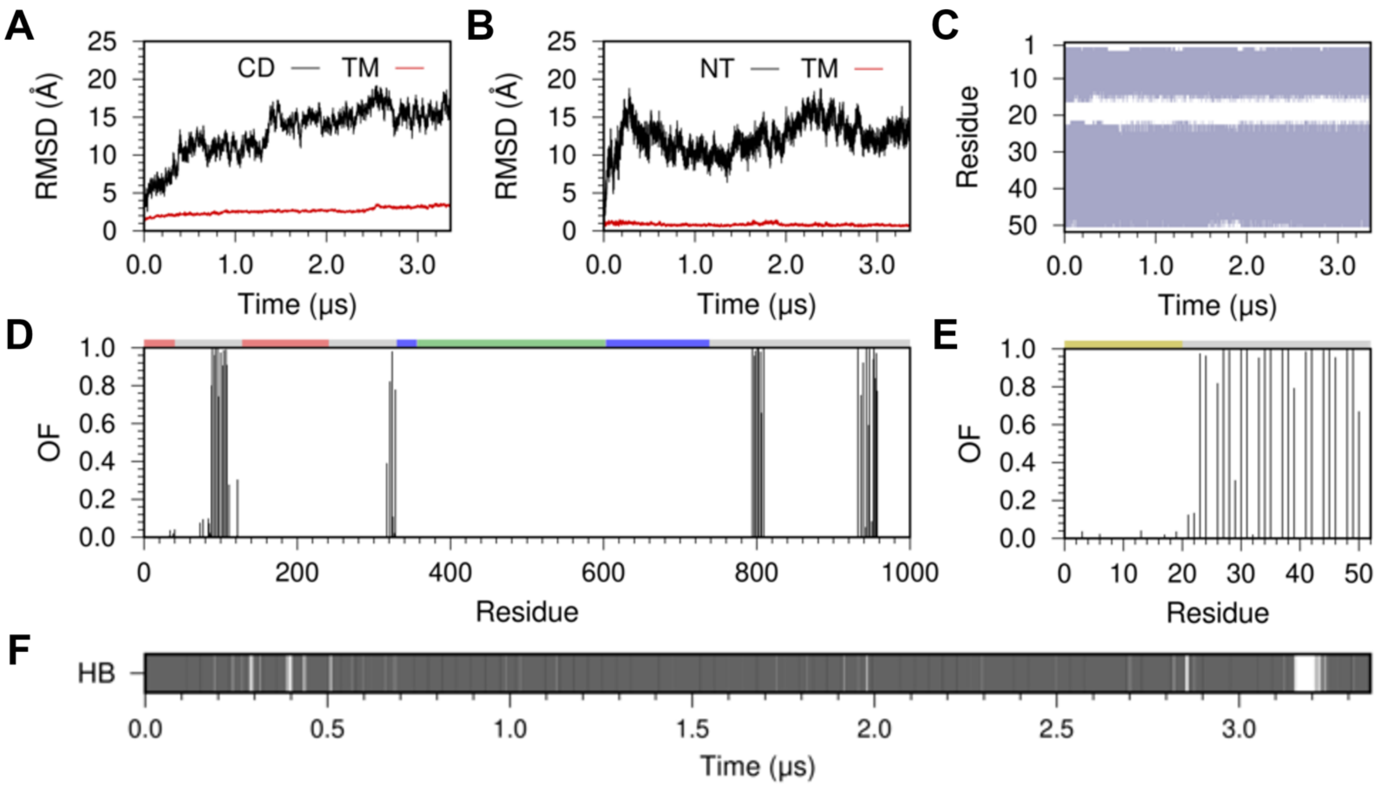


**Fig. S3. Structural analysis of the $Ca^{2+}$-free SERCA–PLB from the MD replicate #3**. (A) The RMSD of the cytosolic (CD, black) and transmembrane (TM, red) domains of SERCA. (B) The RMSD of the cytosolic N-terminal (NT, black) and transmembrane (TM, red) regions of PLB in the complex. In all cases, the RMSD was calculated by aligning the backbone of the TM domain of SERCA with the structure at the beginning of the MD replicate. (C) Secondary structure evolution of PLB in the complex; here, α-helix and random coil are colored as violet and white, respectively. This plot shows that the structure of PLB remains intact in the μs timescale. (D) Occupancy fraction (OF) of PLB in the canonical binding site of SERCA calculated from the MD replicate. This plot indicates that PLB interacts with the TM domain of SERCA (gray) and does not form favorable interactions with the A (red), P (blue) or N (green) domains of SERCA. (E) Occupancy fraction (OF) of each individual residue of PLB in the canonical site. This plot demonstrates that the TM (gray), but not the cytosolic (yellow) domain of PLB interacts with SERCA in the MD trajectory. (F) Time-dependent evolution of the hydrogen bond between PLB residue Asn34 and SERCA residue Gly801. High stability of this interaction is key for SERCA inhibition by PLB [1, 5, 23], thus demonstrating that in the absence of $Ca^{2+}$ SERCA is primarily in an inhibited state.

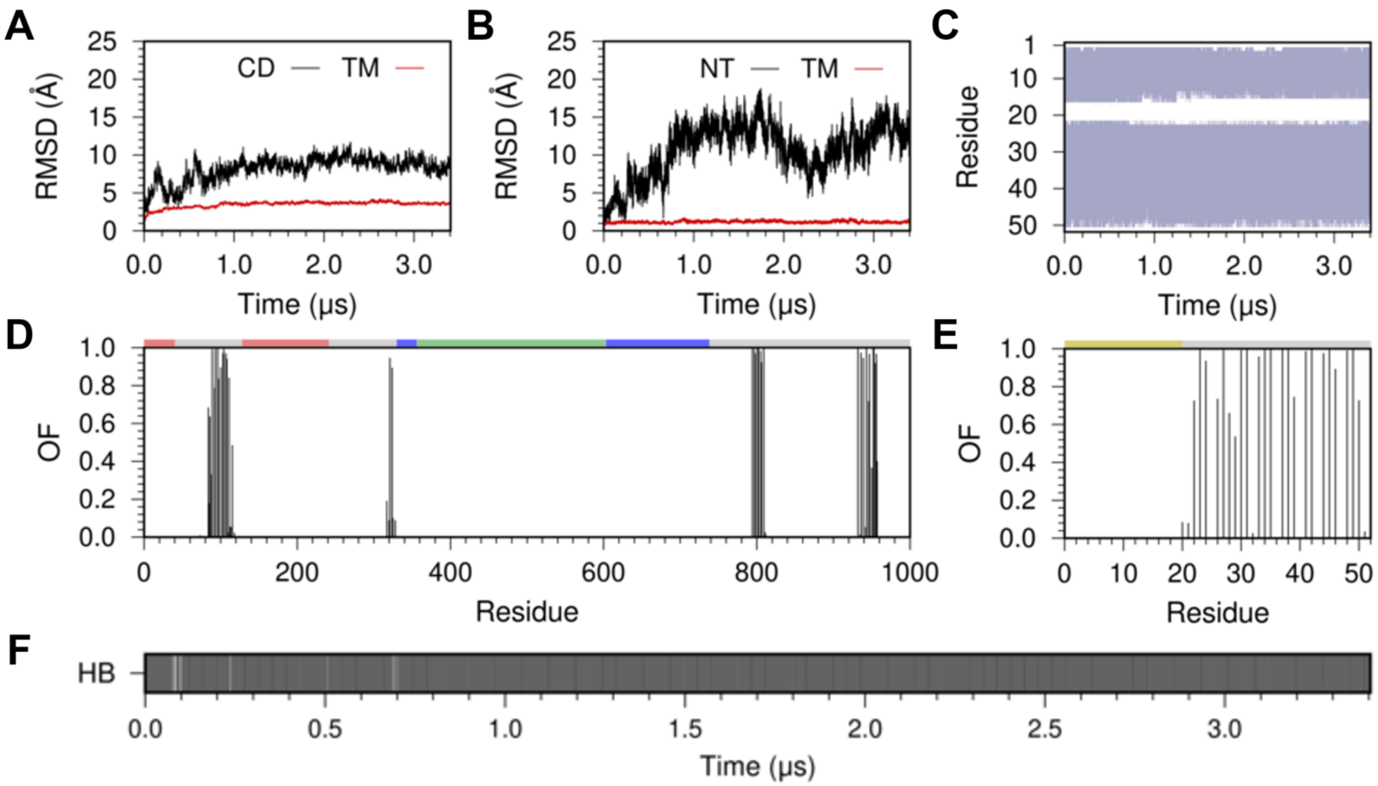


**Fig. S4. Structural analysis of the $Ca^{2+}$-free SERCA–PLB from the MD replicate #4**. (A) The RMSD of the cytosolic (CD, black) and transmembrane (TM, red) domains of SERCA. (B) The RMSD of the cytosolic N-terminal (NT, black) and transmembrane (TM, red) regions of PLB in the complex. In all cases, the RMSD was calculated by aligning the backbone of the TM domain of SERCA with the structure at the beginning of the MD replicate. (C) Secondary structure evolution of PLB in the complex; here, α-helix and random coil are colored as violet and white, respectively. This plot shows that the structure of PLB remains intact in the μs timescale. (D) Occupancy fraction (OF) of PLB in the canonical binding site of SERCA calculated from the MD replicate. This plot indicates that PLB interacts primarily with the TM domain of SERCA (gray) and does not form favorable interactions with the A (red), P (blue) or N (green) domains of SERCA. (E) Occupancy fraction (OF) of each individual residue of PLB in the canonical site. This plot demonstrates that the TM (gray), but not the cytosolic (yellow) domain of PLB interacts with SERCA in the MD trajectory. (F) Time-dependent evolution of the hydrogen bond between PLB residue Asn34 and SERCA residue Gly801. Stability of this interaction is key for SERCA inhibition by PLB [1, 5, 23], thus demonstrating that in the absence of $Ca^{2+}$ SERCA is primarily in an inhibited state.

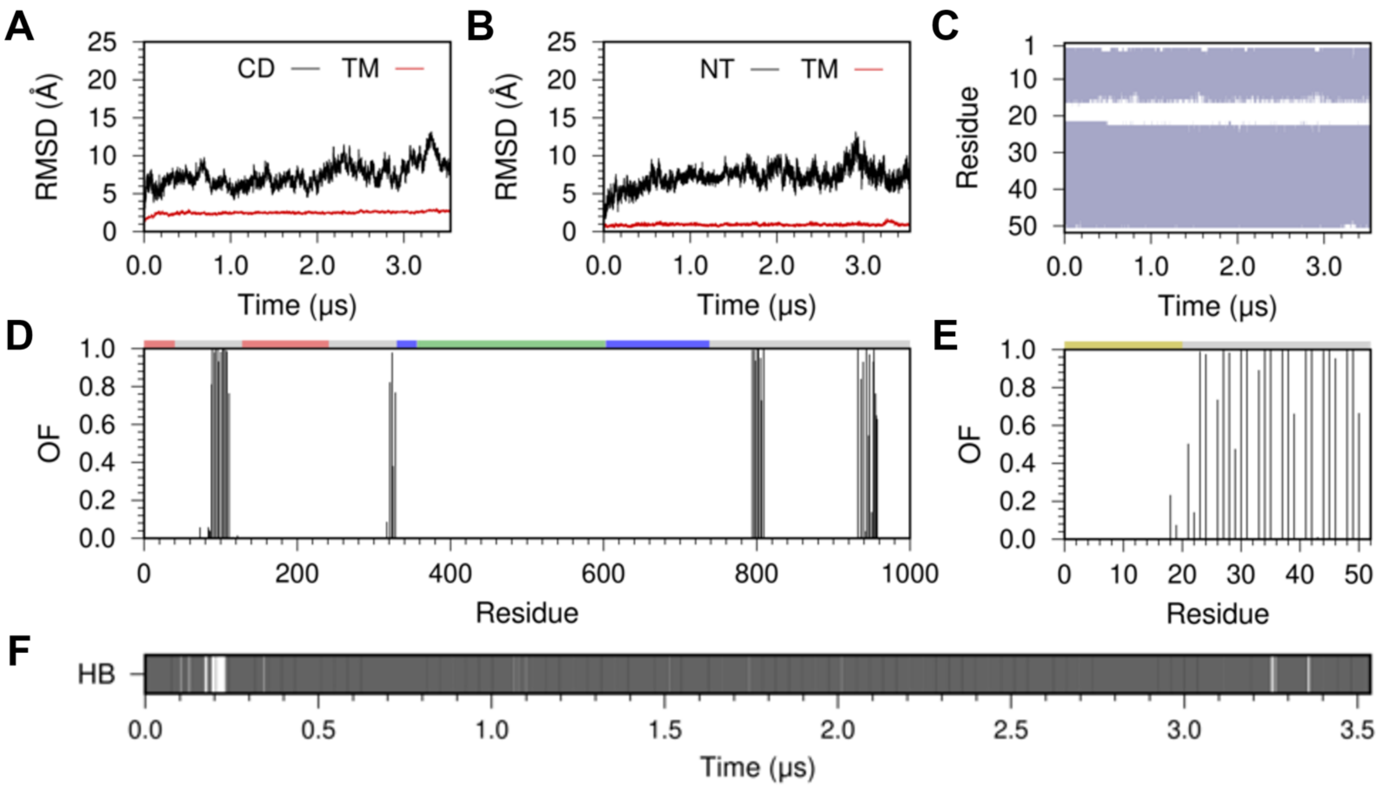


**Fig. S5. Structural analysis of the $Ca^{2+}$-free SERCA–PLB from the MD replicate #5**. (A) The RMSD of the cytosolic (CD, black) and transmembrane (TM, red) domains of SERCA. (B) The RMSD of the cytosolic N-terminal (NT, black) and transmembrane (TM, red) regions of PLB in the complex. In all cases, the RMSD was calculated by aligning the backbone of the TM domain of SERCA with the structure at the beginning of the MD replicate. (C) Secondary structure evolution of PLB in the complex; here, α-helix and random coil are colored as violet and white, respectively. This plot shows that the structure of PLB remains intact in the μs timescale. (D) Occupancy fraction (OF) of PLB in the canonical binding site of SERCA calculated from the MD replicate. This plot indicates that PLB interacts primarily with the TM domain of SERCA (gray) and does not form favorable interactions with the A (red), P (blue) or N (green) domains of SERCA. (E) Occupancy fraction (OF) of each individual residue of PLB in the canonical site. This plot demonstrates that the TM (gray) domain of PLB exclusively interacts with SERCA in the MD trajectory. (F) Time-dependent evolution of the hydrogen bond between PLB residue Asn34 and SERCA residue Gly801. High stability of this interaction is key for SERCA inhibition by PLB [1, 5, 23], thus demonstrating that in the absence of $Ca^{2+}$ SERCA is primarily in an inhibited state.

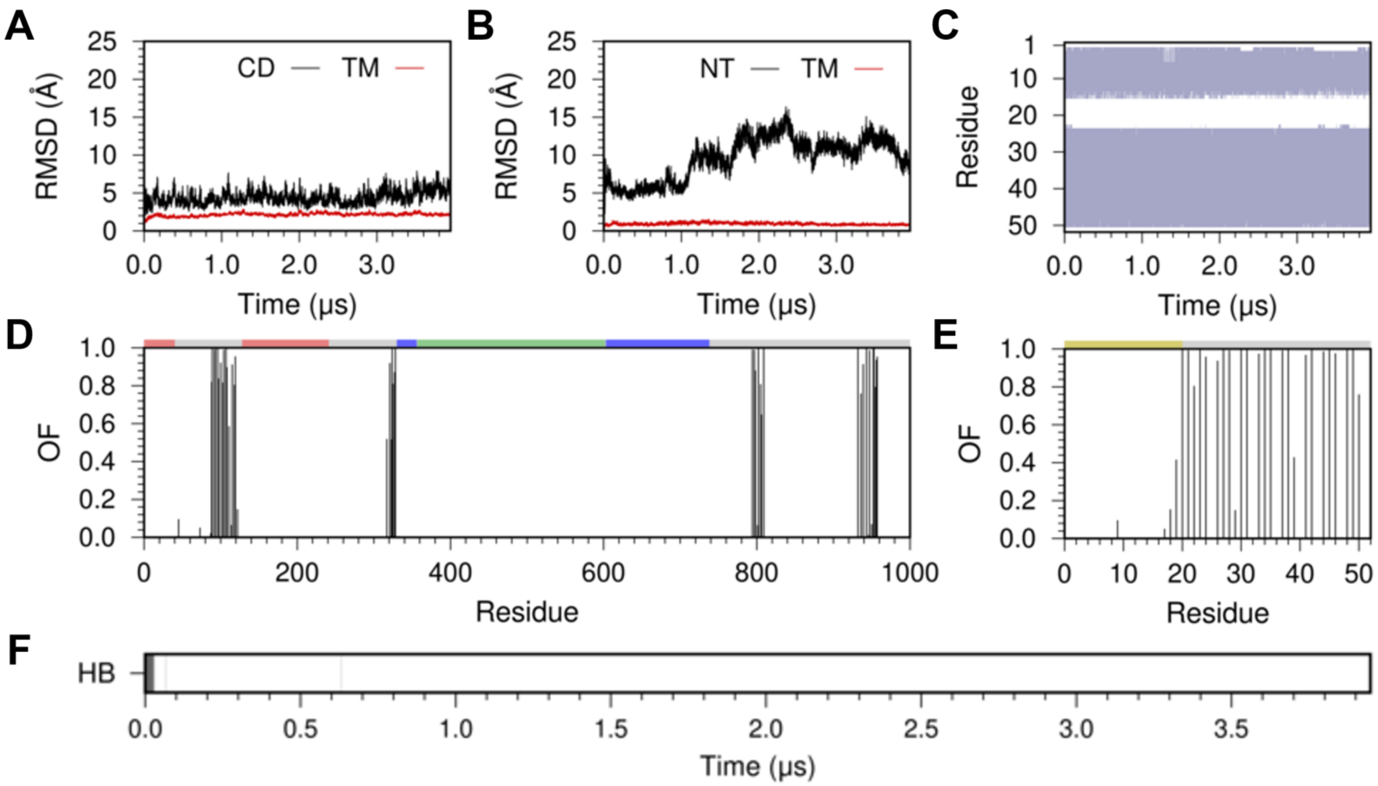


**Fig. S6. Structural analysis of the $Ca^{2+}$-bound SERCA–PLB from the MD replicate #1**. (A) The RMSD of the cytosolic (CD, black) and transmembrane (TM, red) domains of SERCA. (B) The RMSD of the cytosolic N-terminal (NT, black) and transmembrane (TM, red) regions of PLB in the complex. In all cases, the RMSD was calculated by aligning the backbone of the TM domain of SERCA with the structure at the beginning of the MD replicate. (C) Secondary structure evolution of PLB in the complex; here, α-helix and random coil are colored as violet and white, respectively. This plot shows that the structure of PLB remains intact in the μs timescale, thus supporting the notion that $Ca^{2+}$ binding does not induce structural changes in the structural dynamics of PLB.[24] (D) Occupancy fraction (OF) of PLB in the canonical binding site of SERCA calculated from the MD replicate. This plot indicates that in the presence of bound $Ca^{2+}$, PLB interacts primarily with the TM domain of SERCA (gray) and does not form favorable interactions with the A (red), P (blue) or N (green) domains of SERCA. (E) Occupancy fraction (OF) of each individual residue of PLB in the canonical site. This plot demonstrates that the TM (gray) domain of PLB exclusively interacts with SERCA in the MD trajectory of $Ca^{2+}$-bound SERCA–PLB. (F) Time-dependent stability of the hydrogen bond between PLB residue Asn34 and SERCA residue Gly801. The disruption of this interaction in the trajectory demonstrates that binding of a single $Ca^{2+}$ ion relieves key inhibitory interactions between SERCA and PLB [1, 5, 23].

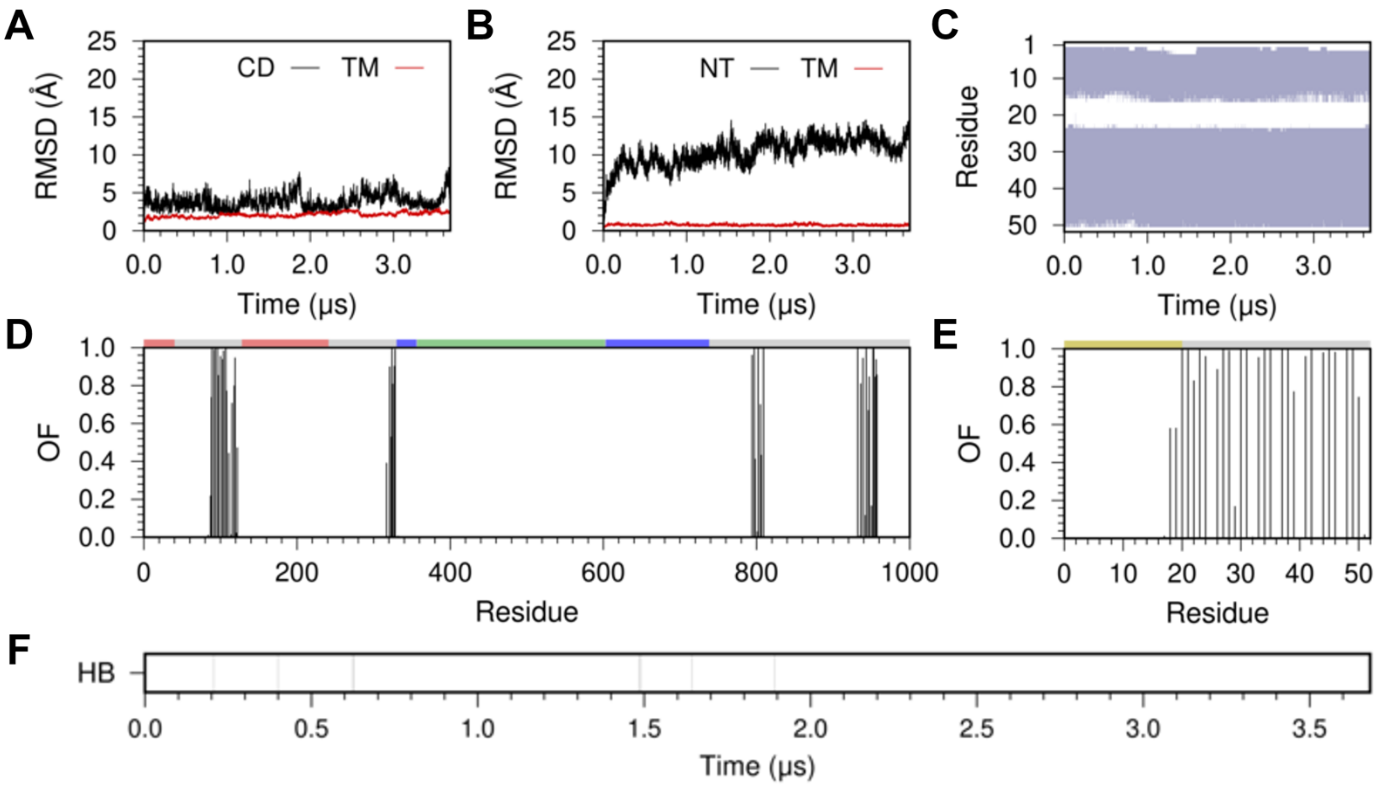


**Fig. S7. Structural analysis of the $Ca^{2+}$-bound SERCA–PLB from the MD replicate #2**. (A) The RMSD of the cytosolic (CD, black) and transmembrane (TM, red) domains of SERCA. (B) The RMSD of the cytosolic N-terminal (NT, black) and transmembrane (TM, red) regions of PLB in the complex. In all cases, the RMSD was calculated by aligning the backbone of the TM domain of SERCA with the structure at the beginning of the MD replicate. (C) Secondary structure evolution of PLB in the complex; here, α-helix and random coil are colored as violet and white, respectively. This plot shows that the structure of PLB remains intact in the μs timescale, thus supporting the notion that $Ca^{2+}$ binding does not induce structural changes in the structural dynamics of PLB.[24] (D) Occupancy fraction (OF) of PLB in the canonical binding site of SERCA calculated from the MD replicate. This plot indicates that in the presence of bound $Ca^{2+}$, PLB interacts primarily with the TM domain of SERCA (gray) and does not form favorable interactions with the A (red), P (blue) or N (green) domains of SERCA. (E) Occupancy fraction (OF) of each individual residue of PLB in the canonical site. This plot demonstrates that the TM (gray), and some residues in the cytosolic extension (residues Ile18 and Glu19) domain of PLB interacts with SERCA in the MD trajectory. (F) Time-dependent stability of the hydrogen bond between PLB residue Asn34 and SERCA residue Gly801. Complete disruption of this interaction in the trajectory demonstrates that binding of a single $Ca^{2+}$ ion relieves key inhibitory interactions between SERCA and PLB [1, 5, 23].

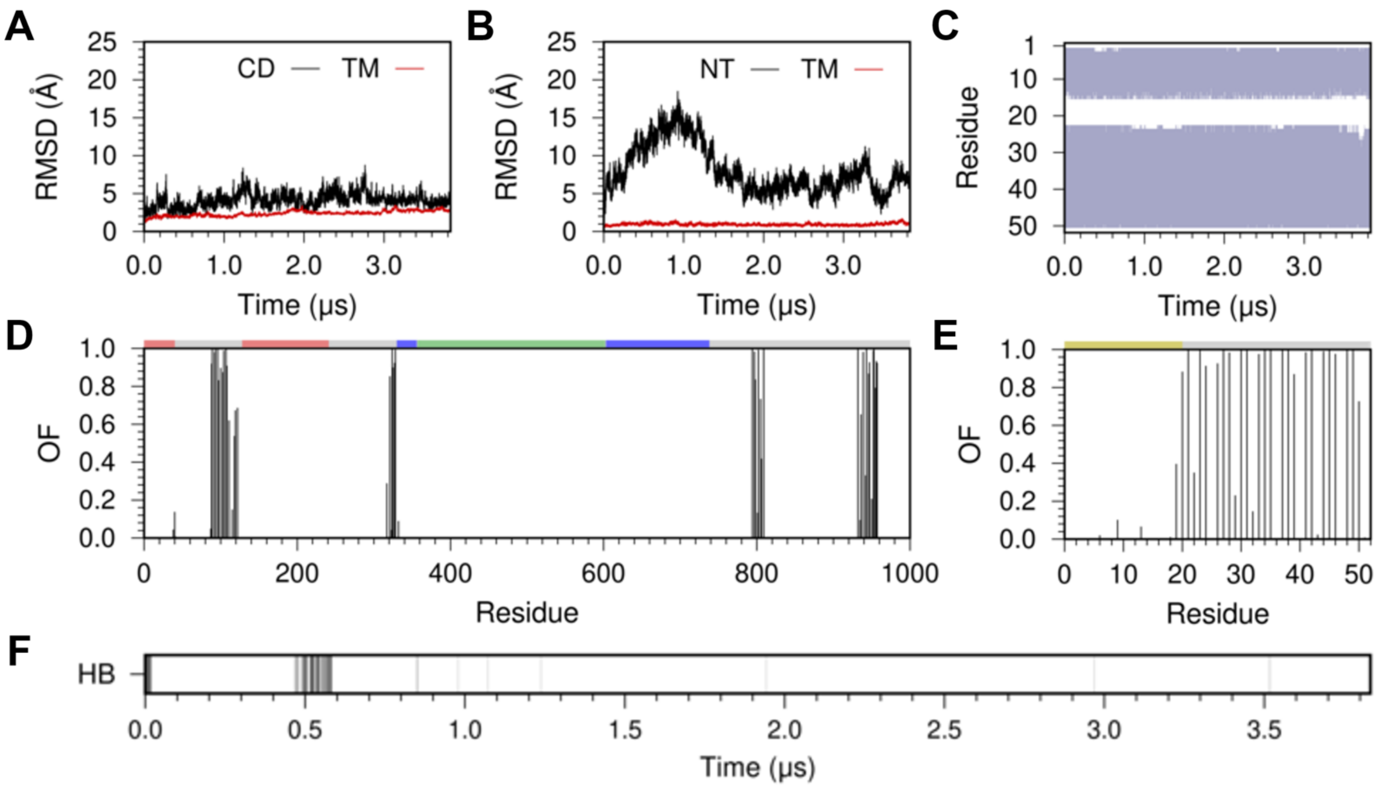


**Fig. S8. Structural analysis of the $Ca^{2+}$-bound SERCA–PLB from the MD replicate #3**. (A) The RMSD of the cytosolic (CD, black) and transmembrane (TM, red) domains of SERCA. (B) The RMSD of the cytosolic N-terminal (NT, black) and transmembrane (TM, red) regions of PLB in the complex. In all cases, the RMSD was calculated by aligning the backbone of the TM domain of SERCA with the structure at the beginning of the MD replicate. (C) Secondary structure evolution of PLB in the complex; here, α-helix and random coil are colored as violet and white, respectively. This plot shows that the structure of PLB remains intact in the μs timescale, thus supporting the notion that $Ca^{2+}$ binding does not induce structural changes in the structural dynamics of PLB.[24] (D) Occupancy fraction (OF) of PLB in the canonical binding site of SERCA calculated from the MD replicate. This plot indicates that in the presence of bound $Ca^{2+}$, PLB interacts primarily with the TM domain of SERCA (gray) and does not form favorable interactions with the A (red), P (blue) or N (green) domains of SERCA. (E) Occupancy fraction (OF) of each individual residue of PLB in the canonical site. This plot demonstrates that the TM (gray) domain of PLB primarily interacts with SERCA in the MD trajectory of $Ca^{2+}$-bound SERCA–PLB. (F) Time-dependent stability of the hydrogen bond between PLB residue Asn34 and SERCA residue Gly801. The disruption of this interaction in the trajectory demonstrates that binding of a single $Ca^{2+}$ ion relieves key inhibitory interactions between SERCA and PLB [1, 5, 23].

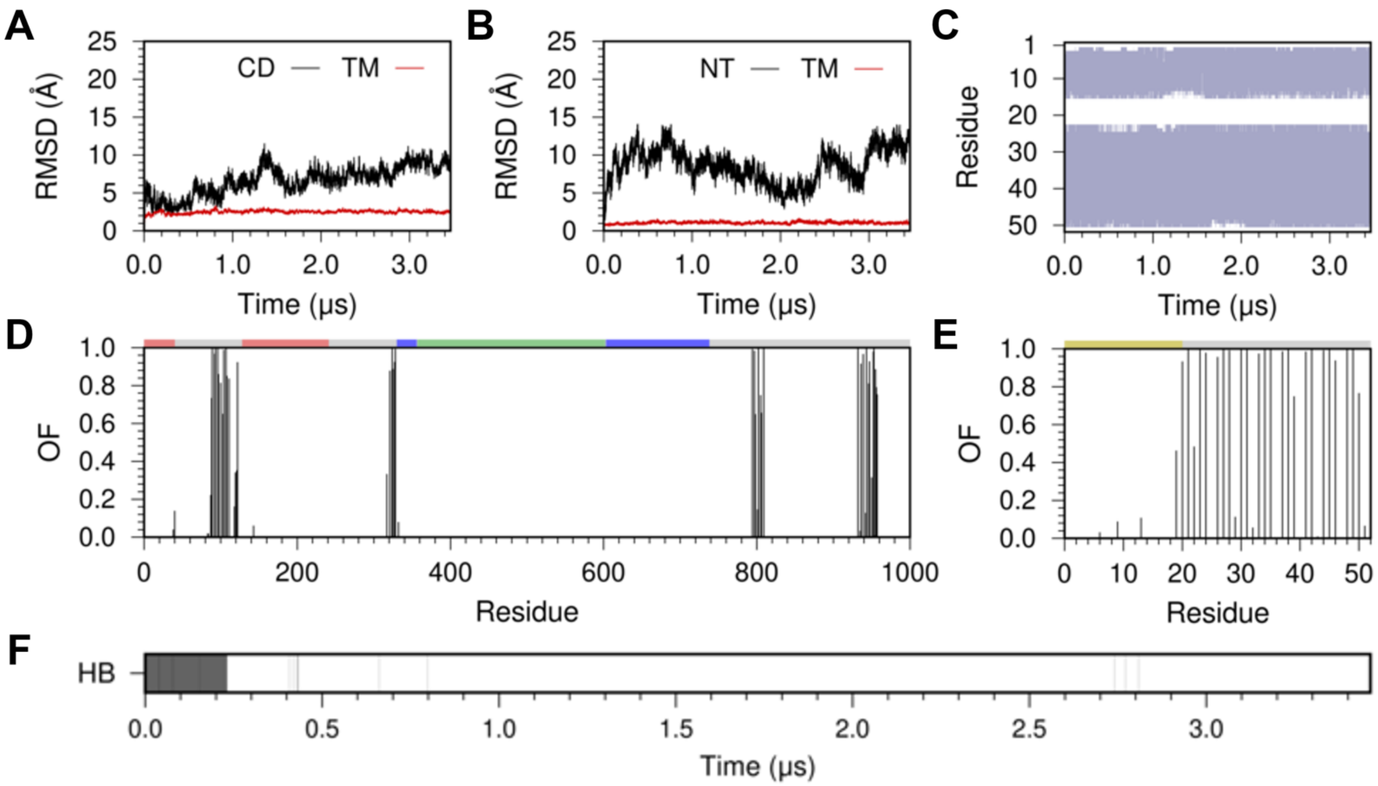


**Fig. S9. Structural analysis of the $Ca^{2+}$-bound SERCA–PLB from the MD replicate #4.** (A) The RMSD of the cytosolic (CD, black) and transmembrane (TM, red) domains of SERCA. (B) The RMSD of the cytosolic N-terminal (NT, black) and transmembrane (TM, red) regions of PLB in the complex. In all cases, the RMSD was calculated by aligning the backbone of the TM domain of SERCA with the structure at the beginning of the MD replicate. (C) Secondary structure evolution of PLB in the complex; here, α-helix and random coil are colored as violet and white, respectively. This plot shows that the structure of PLB remains intact in the μs timescale, thus supporting the notion that $Ca^{2+}$ binding does not induce structural changes in the structural dynamics of PLB.[24] (D) Occupancy fraction (OF) of PLB in the canonical binding site of SERCA calculated from the MD replicate. This plot indicates that in the presence of bound $Ca^{2+}$, PLB interacts primarily with the TM domain of SERCA (gray) and does not form favorable interactions with the A (red), P (blue) or N (green) domains of SERCA. (E) Occupancy fraction (OF) of each individual residue of PLB in the canonical site. This plot demonstrates that the TM (gray) domain of PLB primarily interacts with SERCA in the MD trajectory of $Ca^{2+}$-bound SERCA–PLB. (F) Time-dependent stability of the hydrogen bond between PLB residue Asn34 and SERCA residue Gly801. The disruption of this interaction in the trajectory demonstrates that binding of a single $Ca^{2+}$ ion relieves key inhibitory interactions between SERCA and PLB [1, 5, 23].

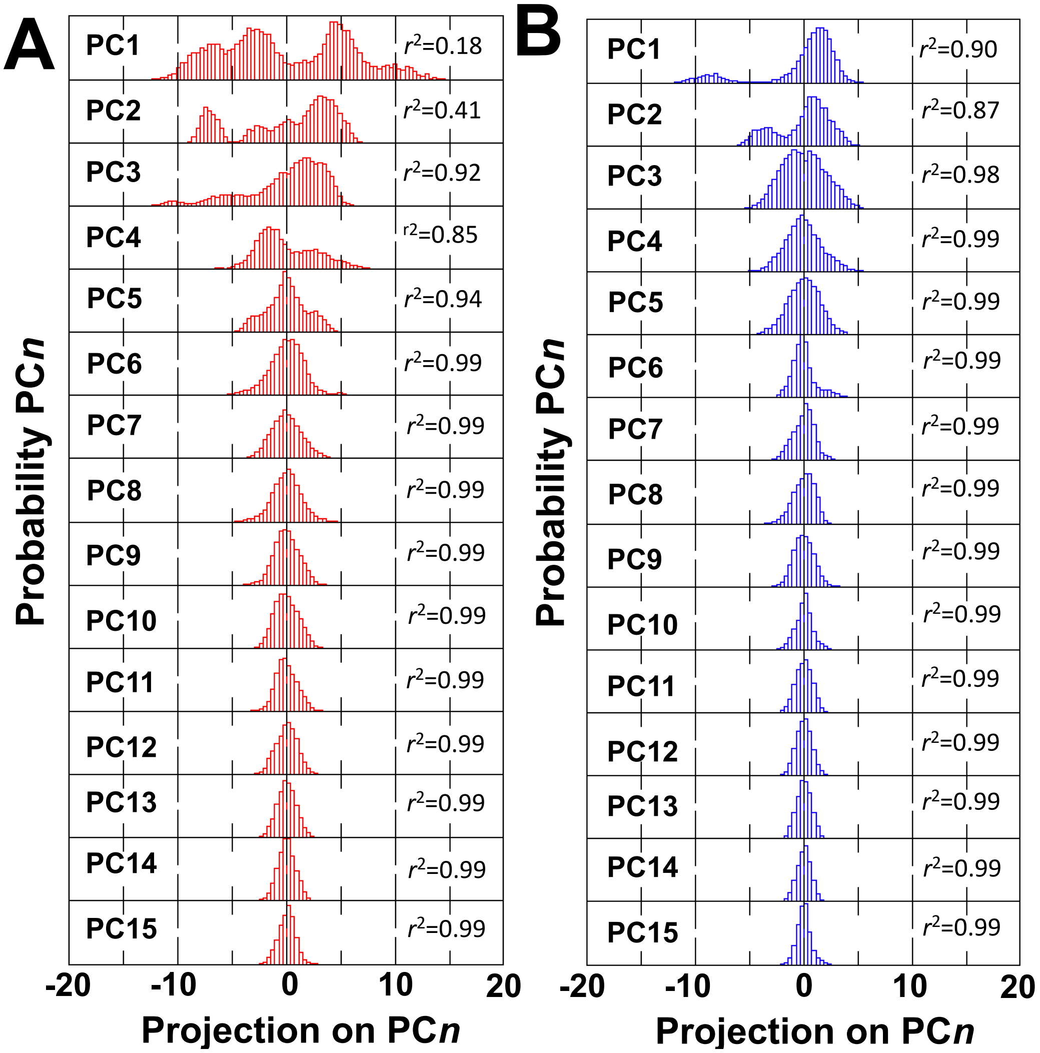


**Fig. S10. Probability distributions for the first 15 principal components (PC) extracted in the trajectories of (A) $Ca^{2+}$-free and (B) $Ca^{2+}$-bound SERCA–PLB complex**. The $r^2$ values shown inside each plot corresponds to the coefficient of determination obtained after fitting each histogram to a one-Gaussian distribution by least squares analysis. Distributions with a coefficient of determination $r^2$ <0.9 is considered to be non-Gaussian (e.g., PC1), and thus belong to the essential phase space [25]. Distributions with $r^2$ values between 0.9 and 0.98 (e.g., PCs 2-5 for the $Ca^{2+}$-free complex) indicate that the principal component retains some significant non-Gaussian features, therefore contribute to the essential phase space. Finally, principal component distributions with $r^2$>0.98 (e.g., PC 6 through 15) are Gaussian fluctuations, thus not considered part of the essential phase space [25].

**Table S1.** Correlation coefficient (*r*) and Pearson's test ($\chi^2$) values computed for the Gaussian fitting models applied to the distance distribution between residues Thr171 and Glu486 obtained from the trajectories of $Ca^{2+}$-free and $Ca^{2+}$-bound SERCA–PLB.

| | SERCA–PLB, without bound $Ca^{2+}$ | | SERCA–PLB, with bound $Ca^{2+}$ | |
|---|---|---|---|---|
| **Model** | **Correlation coefficient (*r*)** | **Pearson's test ($\chi^2$)** | **Correlation coefficient (*r*)** | **Pearson's test ($\chi^2$)** |
| **1-Gaussian** | 0.92 | 0.0335 | 0.98 | 0.0044 |
| **2-Gaussian** | 0.96 | 0.0181 | 0.99 | 0.0042 |
| **3-Gaussian** | 0.97 | 0.0136 | 0.99 | 0.0009 |
| **4-Gaussian** | 0.99 | 0.0005 | – | – |

**Table S2.** Correlation coefficient (*r*) and Pearson's test ($\chi^2$) values computed for the Gaussian fitting models applied to the distance distribution between residues Arg139 and Asp426 obtained from the trajectories of $Ca^{2+}$-free and $Ca^{2+}$-bound SERCA–PLB.

| | SERCA–PLB, without bound $Ca^{2+}$ | | SERCA–PLB, with bound $Ca^{2+}$ | |
|---|---|---|---|---|
| **Model** | **Correlation coefficient (*r*)** | **Pearson's test ($\chi^2$)** | **Correlation coefficient (*r*)** | **Pearson's test ($\chi^2$)** |
| **1-Gaussian** | 0.98 | 0.0028 | 0.98 | 0.0089 |
| **2-Gaussian** | 0.99 | 0.0007 | 0.99 | 0.0011 |
